# Sudan black lipid blot – a rapid and simple method for quantification of lipids in biological samples

**DOI:** 10.1101/2022.07.27.501748

**Authors:** Jan Homolak, Mihovil Joja, Pavel Markovic, Melita Salkovic-Petrisic

**Affiliations:** Department of Pharmacology, University of Zagreb School of Medicine, Zagreb, Croatia; Croatian Institute for Brain Research, University of Zagreb School of Medicine, Zagreb, Croatia

## Abstract

Bioanalytical techniques for the isolation and quantification of total lipids in biological samples are an integral part of lipidomic workflows and widely used tools for metabolic assessment at the cellular and organismic levels. The most widely used protocol for the isolation, extraction, and quantification of total lipids in biological tissues was originally introduced by Folch et al.. It requires a relatively large amount of tissue and large volumes of lipid extracts for reliable assessment of lipid content using the gravimetric technique. Here, we propose a new method to overcome the aforementioned challenges based on the hypothesis that the partitioning coefficient of the widely used lysochrome diazo dye Sudan Black B between the lipid extract and ethylene glycol can be used to indirectly estimate the absolute concentration of lipids. The proposed method demonstrates great precision and linearity, requires minimal equipment, and enables the analysis of total lipid content in biological specimens available only in limited amounts by reducing the requirements for the input quantity by >300-fold for some tissues (e.g. fecal samples).

## 1. Introduction

Lipidomics is an emerging field of research focused on understanding the structure and function of a set of lipids present in the organism. Tools and methods for studying lipids in different biological samples are becoming increasingly important considering the fact that lipids play diverse physiological roles and that alterations of the lipidome have been implicated in the etiopathogenesis of many medical conditions. Although sophisticated analytical methods often relying on the principles of mass spectrometry are indispensable for a comprehensive qualitative and quantitative overview of the lipidome, simple biochemical techniques for the isolation and quantification of total lipids are still widely used in many laboratories as i) a loading control technique used to enable assessment of relative changes in a particular lipid species by introducing the total lipid content of the tested sample as a covariate in the model; and/or ii) a screening tool used for a rapid assessment of the overall lipid metabolism (e.g. ^[1,2]^). One of the most widely used methods for the direct assessment of tissue total lipid content is the gravimetric analysis of the evaporated lipid chloroform extract first introduced by Folch et al.^[3]^ that has been successfully adapted for the analysis of total lipid content in different biological specimens (e.g.^[4]^). The original method proposed by Folch et al. has its unquestionable advantages: it has minimal requirements in regards to the equipment and reagents, it is cost-efficient, and it relies on a direct assessment of the lipid weight which makes it impervious to chemical bias (e.g. the interference of the sample matrix with the horseradish peroxidase reporter system present in many widely used biological assays). Nevertheless, Folch’s method also has several disadvantages: i) The original method requires a considerable amount of tissue to be analyzed in order to extract lipids in a quantity that can be reliably assessed using a standard laboratory analytical balance (e.g. 1000 mg in the protocol proposed by Kraus et al.^[4]^) ; ii) A large volume of the chloroform extract that has to be obtained from each sample prolongs the analysis time as it can take several days for the liquid to fully evaporate (e.g. 3.3. ml of chloroform in the protocol proposed by Kraus et al. ^[4]^); iii) The evaporation step requires a relatively large area inside a fume hood to be occupied with evaporating tubes/dishes for days; iv) The need for a large quantity of chloroform extract can result in increased exposure of the laboratory personnel to toxic fumes with potential health consequences ^[5]^. Here, we propose a new method to overcome the aforementioned challenges based on the hypothesis that the partitioning coefficient of the widely used lysochrome diazo histological dye Sudan Black B (SBB; (2,2-dimethyl-1,3-dihydroperimidin-6-yl)-(4-phenylazo-1-naphthyl)diazene) between the lipid extract obtained from the biological sample and ethylene glycol (that does not extract lipids but dissolves sufficient quantities of the liposoluble SBB ^[6]^) can be used to estimate the absolute concentration of lipids present in the biological sample.

The aim was to assess i) whether the simple deposition of the chloroform lipid extract on the microscope slide followed by staining in the SBB dissolved in ethylene glycol can be used to reliably assess lipid concentration (using the Folch’s method as the gold standard) in different biological samples; and if latter is the case ii) whether the new protocol can enable a reduction in the amount of lipid extract required for the analysis so that it can be performed using biological specimens available only in limited quantities (at the fraction of the cost).

## 2. Materials and methods

### 2.1. Sample preparation

Fecal samples were collected from 3-month-old male Wistar rats from the animal facility at the Department of Pharmacology (University of Zagreb School of Medicine, Zagreb, Croatia) animal facility. The animals were kept under standard conditions: 3/cage in the controlled environment with 21–23 °C room temperature, 40–70% relative humidity, a 12 h light/dark cycle (7/19CET), standard bedding changed twice per week, and standardized food pellets and water available *ad libitum*. Fecal pellets were separated from bedding material using a sieve and left at room temperature until all the pellets were completely dry. Dry feces was ground using a commercial coffee grinder and fecal powder was stored at room temperature for subsequent use. To assess whether the proposed method was suitable for the analysis of total lipid content in biological specimens other than feces we used samples of rat brain and pancreatic tissue from 5-month-old male Wistar rats used in previous experiments and stored at -80°C in concordance with the 3R principles^[7]^. In the original experiments, the animals were euthanized in deep general anesthesia (ip ketamine 70 mg/kg, xylazine 7 mg/kg) and decapitated, pancreatic and brain tissue was dissected, snap frozen in liquid nitrogen, and stored at -80°C. The animal study from which the tissue was obtained was approved by the University of Zagreb School of Medicine Ethics Committee (380-59-10106-18-111/173) and the Croatian Ministry of Agriculture (EP 186/2018). The brain and the pancreatic tissue samples were placed in the lysis buffer containing 150 mM NaCl, 50 mM Tris-HCl, 1 mM EDTA, 1% Triton X-100, 1% sodium deoxycholate, 0.1% SDS, 1 mM phenylmethylsulfonyl fluoride, protease inhibitor cocktail (Sigma-Aldrich, USA) and PhosSTOP phosphatase inhibitor (Roche, Switzerland) and homogenized with the Microson Ultrasonic Cell Disruptor (Misonix, SAD) on ice.

### 2.2. Isolation, purification, and quantification of total lipids using Folch’s method

A widely used method for the isolation and purification of total lipids from animal tissues originally proposed by Folch et al. ^[3,4]^ was used to estimate the total lipid content of all three biological specimens (feces, brain, and the pancreas). Lipid extraction from feces was done following the modified Folch’s protocol described by Kraus et al. ^[4]^. Briefly, 5 ml of 1xPBS was added to 1 g of fecal powder and vortexed thoroughly. 5 ml of 2:1 (v/v) chloroform in methanol was added, vortexed again, and the suspension was spun down at an RCF of 1000 x g for 10 minutes to obtain two liquid phases separated by a solid phase of insoluble material. The lipid (lower) phase was then isolated and placed in the glass Petri dish previously weighed on an analytical balance. The glass dish containing the lipid extract was placed under a fume hood until all liquid was evaporated. The empty dish weight was then subtracted from the weight of the same dish containing unevaporated material to obtain the weight of lipids in milligrams and to estimate the concentration of lipids in the aliquot of the same sample. The same protocol was used for the extraction of lipids from the brain and the pancreatic tissue homogenates. The whole procedure was repeated until the cumulative weight of the extracted lipids from each tissue was large enough (>5 mg) so that it can be reliably assessed using the analytical balance (Mettler Toledo, Columbus, Ohio, SAD). When the optimal weight (for fecal powder) or volume (for the brain and the pancreatic tissue homogenates) was determined for each tissue, the whole procedure was repeated with the required volume (i.e. number of separate conical centrifuge tubes) processed in parallel to obtain the volume of sample that can be reliably measured using the Folch’s procedure and the novel method proposed here. The repeated isolation was conducted to avoid bias that may have been introduced by evaporation and/or chemical alteration of the lipid extract aliquot that was used as a standard sample to generate the SBLB calibration curve.

### 2.3. Sudan black lipid blot (SBLB)

The proposed protocol (Sudan black lipid blot; SBLB) was developed as an alternative way to estimate lipid concentration in the chloroform layer following the 2:1 chloroform in methanol lipid extraction. In the protocol proposed here, 1 _μ_l of the lipid-containing chloroform extract is pipetted onto a standard microscope slide placed on top of a laboratory heater (pre-heated to 40 °C) under a fume hood. A small volume and heated surface enable rapid evaporation of chloroform and fixation of residual contents onto the glass slide. After all samples are administered, the slide is placed in the SBB staining solution containing 1% SBB dissolved in ethylene glycol for 20 s and destained under running H_2_O to remove excess dye from the slide. Commonly used SBB alcohol solvents (e.g. ethanol) are not compatible with the proposed method as the solvent of choice should prevent the loss of lipid particles from the slide and be completely inert with respect to the solubility of lipid material in it ^[6,8]^. After destaining, the slide is air dried at room temperature and digitalized using the office scanner or camera. The image/scan of the microscope slide is then analyzed using a standard procedure for dot blot quantification (e.g. as described in detail for nitrocellulose redox permanganometry^[9]^). The concentration of lipids is determined from the standard concentration curve by quantifying integrated density for each sample based on the assumption that signal density reflects the partitioning coefficient between the solvent (i.e. ethylene glycol) and the lipid extract.

For SBLB to provide the most reliable results a single sample with known lipid concentration should be used to generate a serial dilution for the determination of a standard curve. In order to avoid bias that may be introduced by possible qualitative and quantitative differences in lipid content between different tissues, we propose a fresh lipid extract from the same tissue to be analyzed as a standard. Considering that Folch’s protocol provides a direct and reliable way to assess absolute lipid concentration we propose that, before SBLB, a single sample is analyzed using the standard Folch’s procedure. The aliquot of the evaporated sample can then be used for the generation of the calibration curve for SBLB, while the assessment of linearity and coefficient of variation (CV) at different target concentrations can be used as quality control to ensure analytical validity of the assay in the tissue of interest. Finally, as SBLB is compatible with parallel processing of many samples, we suggest that both standard and test samples are measured in multiple technical replicates.

## 3. Results

### 3.1. SBLB is a time-efficient and reliable method for the assessment of fecal lipids

An overview of the proposed SBLB method is depicted in Fig 1A,B. Briefly, for quantitative assessment of total lipid content in fecal samples, a single sample is first processed using the standard Folch’s procedure for direct assessment of absolute lipid concentration. The aliquot of the chloroform lipid extract of the standard fecal sample is used to generate serial dilutions for the calibration curve processed in parallel with other samples (Fig 1A). Fecal samples to be analyzed using the SBLB are first subjected to a standard lipid extraction protocol: 100 mg of fecal powder is dissolved in 500 _μ_l 1xPBS and then mixed with 500 _μ_l of 2:1 (v/v) chloroform in methanol. The samples are centrifuged at an RCF of 1000 x g for 10 minutes to obtain two liquid phases separated by a solid phase of insoluble material and the lower phase (i.e. the chloroform lipid extract) is transferred to another tube and stored. A microscope slide is placed onto the laboratory heater set at 40°C inside a fume hood and 1 μl of each sample (including the samples used for the generation of the standard curve) is pipetted vertically onto the slide in a predetermined order (Fig 1B). A helpful trick is to use a template to mark the position of each sample on the backside of the glass slide with a marker that can be removed with 70% ethanol before the slide is immersed into the SBB staining solution. Once all samples are completely dry the slide is placed in the staining solution (1% SBB in ethylene glycol) for 20 seconds (Fig 1B). The slide is removed from the staining solution and placed under running water to remove excess (unbound) dye (Fig 1B), dried at room temperature, digitalized, and analyzed following a standard protocol for dot blot quantification (Fig 1B).

**Fig 1.**
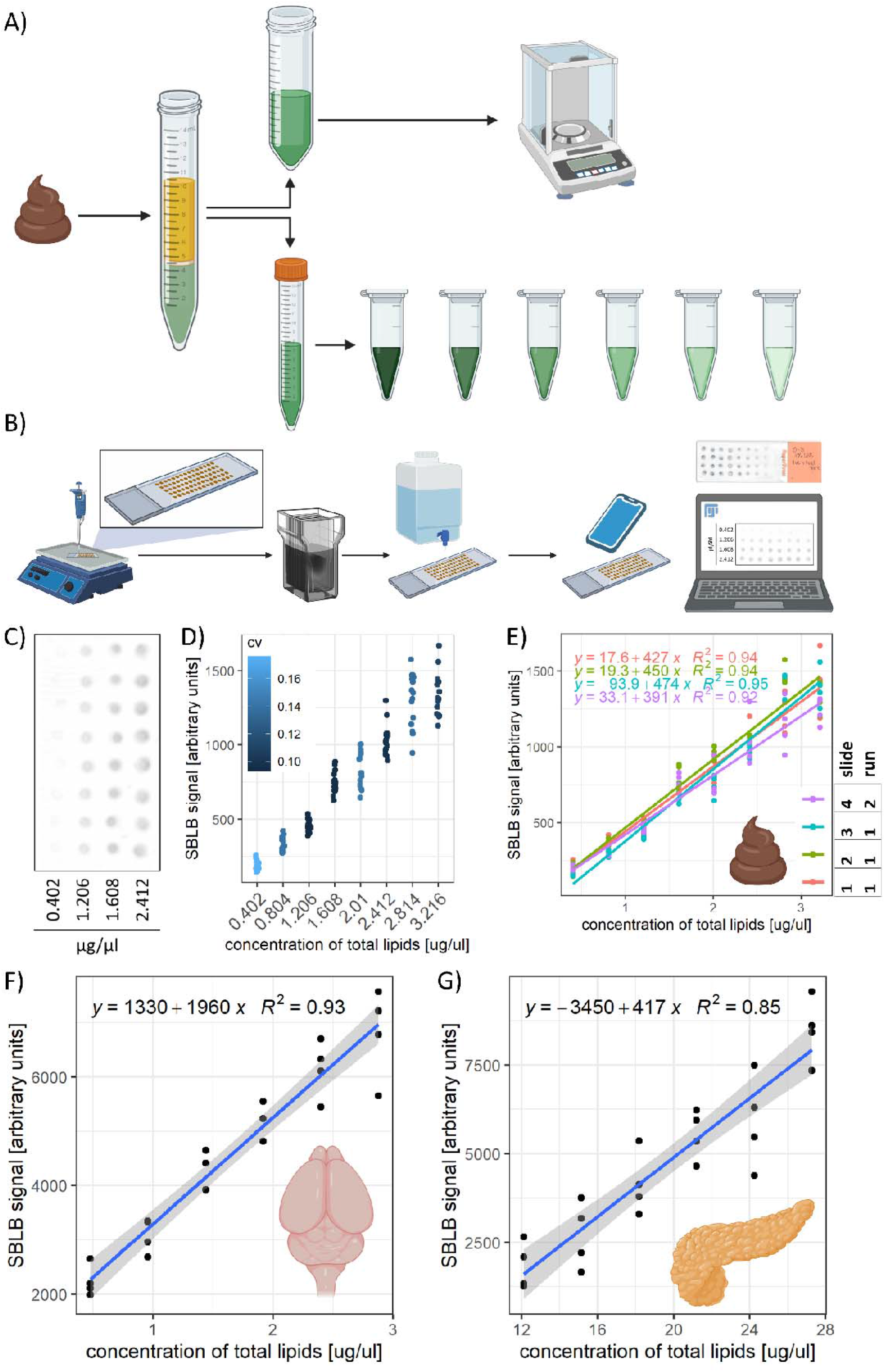
An overview of the Sudan black lipid blot (SBLB) method. (A) A schematic representation of the sample preparation procedure used in the validation experiments. Lipids were extracted from the ground 24-hour rat fecal samples using the standard procedure of extraction in the 2:1 (v/v) chloroform in methanol solution introduced by Folch et al. ^[3]^. An aliquot of the chloroform fraction containing lipids was used for the preparation of serial dilution while the rest was evaporated under a fume hood in a beaker previously weighed on the analytical balance. The concentration of lipids in the aliquot was assessed by dividing the mass of total lipids left in the Petri dish after solvent evaporation and the total volume of the chloroform extract before the evaporation. (B) A schematic representation of the SBLB method. Briefly, the samples are pipetted onto the pre-marked microscope glass slide placed on top of the laboratory heater inside a fume hood in a predetermined order. The marker is removed from the backside of the slide with a paper towel dipped in 70% ethanol and the slide is placed in a Coplin jar filled with the Sudan black B (SBB) staining solution (1% SBB in ethylene glycol) for 20 seconds. The excess dye is removed from the slide by brief washing under running water and the slide is left to dry out completely at room temperature. The slide is digitalized using a scanner or camera and lipid content is estimated using a standard procedure for dot blot quantification. An example of an SBLB microscope slide is shown. (C) A representative scan of the microscope slide obtained by the SBLB protocol. (D) The SBLB bioanalytical method validation demonstrates good linearity and precision in the assessment protocol testing within-run parallel processing (3 microscope slides) and between-run drift (1 microscope slide on a subsequent day) using 8 serial dilutions of the standard chloroform extract of fecal lipids with absolute lipid concentration confirmed using the Folch’s method. (E) The results from D with separate linear regression models fitted for each within-run and between-run trial demonstrate a minimal within-run error, and only a slight between-run drift (possibly caused by instability of chloroform extracts). (F) The results from the SBLB procedure applied to the lipid extracts obtained from the homogenate of the rat brain show good linearity and precision. (G) The results from the SBLB procedure applied to the lipid extracts obtained from the homogenate of the rat pancreatic tissue show good linearity and precision.

The satisfactory repeatability of the proposed method is evident already upon visual inspection of the SBLB microscope slides following repeated administration of the serial dilution samples (Fig 1C) and confirmed with the bioanalytical method validation procedures. Assessment of the CV at 8 concentration levels between 0.4 and 3.2 _μ_g/_μ_l indicates good precision satisfying the criteria of the European Medicines Agency bioanalytical method validation guidelines (<15% CV for the non-lower limit of quantification quality control samples)(Fig 1D)^[10]^.

To illustrate the flexibility and robustness of SBLB the validation protocol was conducted using several microscope slides analyzed in parallel during the same run (to estimate the error introduced by the microscope slide – a composite of the effects of staining and destaining both of which are susceptible to human error and through which slides go in sequential order)(Fig 1E). Although we assumed the placement of samples on separate slides would introduce a substantial error and hinder direct within-run between-slide comparisons the results indicate that the potential error introduced by the arrangement of samples on separate slides was minimal and that it would probably not prevent direct comparisons under the assumption that the samples were analyzed in the same run (i.e. under identical heating and pipetting condition, applied by the same researcher, and stained by the same reagents)(Fig 1E). Interestingly, an attempt to analyze the residual composite error introduced by the whole SBLB procedure (i.e. between-run variability assessed by re-analysis of the same samples on separate days and with separate reagents that should approximately reflect the aforementioned variation introduced by heating conditions and human error in the preparation of reagents and pipetting) indicates that even between-run error is relatively modest with samples demonstrating comparable values and good linearity on a subsequent day (Fig 1E). It is important to notice that chloroform extracts are inherently unstable and that the reported between-run error is probably inflated by the effect of sample distortion (e.g. due to chloroform evaporation during sample handling).

### 3.2. SBLB can be used for rapid total lipid quantification in different biological tissues

Finally, to assess whether the SBLB protocol for rapid quantification of total lipids is compatible with different specimens (due to the assumed quantitative and qualitative variability of lipid content across tissues) we analyzed rat brain and pancreatic tissue homogenates following the same procedure and assessed the linearity of the SBLB signal in serial dilutions of the standard samples in which we previously determined total lipid content using the Folch’s method. Although the coefficient of determination was slightly lower (0.93 and 0.85 for the brain and the pancreatic tissue respectively) in comparison with the fit obtained for the fecal sample model, both lipid extracts from the brain and the pancreatic tissue were compatible with the proposed method and demonstrated good linearity across tested concentrations (Fig 1F,G). Since the proposed method seems to be compatible with 3 different tissue samples (feces, the brain, and the pancreatic tissue) we consider it more likely than not that the same or slightly modified procedure can be successfully applied to a wide variety of biological samples. Nevertheless, considering that we can only assume that the cause for greater variability observed in the pancreatic tissue in comparison with fecal samples can be at least partially explained by qualitative and quantitative differences in lipid profiles of analyzed tissues, some caution should be exercised and the proposed method should only be applied if the satisfactory assay performance has been achieved using serial dilutions of the standard sample of the tissue that is to be analyzed.

## 4. Discussion

The presented results provide solid evidence that the SBLB method can be used to reliably estimate the absolute concentration of lipids in different biological samples. Furthermore, as only 1 _μ_l of each sample is required for the SBLB total lipid content analysis, the original Folch’s lipid isolation and extraction protocol can easily be miniaturized. For example, the protocol for lipid extraction from rodent feces ^[4]^ can easily be reduced ∼ 330-fold (requiring only 6.6 _μ_l instead of 3300 _μ_l of chloroform per sample) while still yielding enough extract for several technical replicates for each sample. As a consequence, the exposure of laboratory workers to toxic chloroform can be dramatically reduced. Furthermore, a dramatic reduction in the required volume of lipid extract required for the analysis also means that instead of the usual 1000 mg of powdered feces required for the analysis ^[4]^, total lipid content can be successfully measured in only ∼3 mg of fecal powder. This miniaturization is exceptionally useful for studies in mice where researchers usually had to pool fecal pellets over several days to collect enough samples for the analysis (the mean fecal pellet weight in mice is in the range of several milligrams – e.g. ∼ 6 mg reported by Hoibian et al. ^[11]^). Considering that lipid biosynthesis, transport, and metabolism have all been shown to be under circadian regulation^[12]^ miniaturization of the assay enabled by the introduction of the SBLB method will enable the researchers to assess lipid content in individual pellets, and thus elucidate and/or account for circadian variations in fecal lipid content. Moreover, the SBLB method is exceptionally fast – a batch of 36 samples (the custom-made template we use enables us to deposit 36 samples on a single microscope slide) can easily be analyzed in under 30 minutes. As our validation data (Fig 1D,E) indicates that many slides can be analyzed in parallel, the aforementioned setup can easily be multiplexed and hundreds of lipid extracts could theoretically be analyzed in a single day providing a unique affordable high-throughput platform for the exploration of changes in total lipid content in different biological specimens. Finally, the SBLB method requires minimal reagents and laboratory equipment what makes it easy for implementation in laboratories with limited equipment and/or funding contributing to a reduction in the research capacity gap between the low/middle-income and high-income countries recognized as an important global problem^[13]^.

## 5. Conclusion

The SBLB is a time-efficient and reliable method for the assessment of total lipids in biological samples. As only a small volume (1 _μ_l) of the lipid extract is needed for the analysis of lipid content, the method enables miniaturization of the Folch’s lipid extraction protocol enabling researchers to process many samples in parallel, obtain the results rapidly, minimize the exposure to toxic chloroform, and study lipid content of biological samples that are available only in limited amounts (e.g. mouse fecal pellets).

## 6. Acknowledgments

**JH** conceptualized the method and designed the experimental protocol and validation tests. **JH, MJ**, and **PM** conducted validation experiments. **JH** conducted data analysis and wrote the first draft of the manuscript. **MJ, PM**, and **MSP** provided critical feedback. **MSP** is head of the Laboratory for Molecular Neuropharmacology where all experiments were conducted, principal investigator of the Croatian Science Foundation-funded project (IP-2018-01-8938), and mentor of JH. All authors agreed to the final version of the manuscript.

This work was funded by the Croatian Science Foundation (IP-2018-01-8938). The research was co-financed by the Scientific Centre of Excellence for Basic, Clinical, and Translational Neuroscience (project Experimental and clinical research of hypoxic-ischemic damage in the perinatal and adult brain; GA KK01.1.1.01.0007 funded by the European Union through the European Regional Development Fund).

## 7. Conflict of interest

None.

## Notes

### Competing Interest Statement

The authors have declared no competing interest.

